# The central clock drives metabolic rhythms in muscle stem cells

**DOI:** 10.1101/2025.01.15.633124

**Authors:** Valentina Sica, Jacob G Smith, Oleg Deryagin, Eva Andres, Vera Lukesova, Mirijam Egg, Nina Cabezas-Wallscheid, Salvador Aznar Benitah, Antonio L. Serrano, Eusebio Perdiguero, Pura Muñoz-Cánoves

**Affiliations:** Universitat Pompeu Fabra (UPF), Department of Medicine and Life Sciences (MELIS), 08003 Barcelona, Spain; Department of Cell Biology, Physiology and Immunology, School of Biology, University of Barcelona, Barcelona, Spain; Institute of Biomedicine of the University of Barcelona (IBUB), Barcelona, Spain; Max Planck Institute of Immunobiology and Epigenetics; Institute for Research in Biomedicine (IRB), Barcelona, The Barcelona Institute of Science and Technology, 08028 Barcelona, Spain; Catalan Institution for Research and Advanced Studies (ICREA), 08010, Barcelona, Spain; Altos Labs Inc, San Diego Institute of Science, San Diego, CA 92121, USA

**Keywords:** circadian rhythms, inter-organ crosstalk, stem cells, satellite cells, quiescence, metabolism, autophagy

## Abstract

Circadian rhythms are essential for organismal health. Satellite cells (SCs), the muscle resident stem cells, maintain a state of quiescence yet exhibit robust circadian oscillations at the transcriptional level. Although peripheral clocks have been extensively studied in various tissues, how the intrinsic clock of stem cells interacts with the central, distal clock is largely unknown. We used SC-specific reconstitution of the essential clock gene *Bmal1* to elucidate the role of the local SC clock and its interplay with the central clock in the mouse brain and found that daily transcriptional control of metabolic processes in SCs depend on central clock input, independent of the SC clock. Central clock-driven genes were involved in lipid metabolism, functionally important for SC-mediated muscle repair, and autophagy was required for their oscillation. In summary, we provide the first evidence of circadian coordination of central and local clocks for control of rhythmic gene expression in quiescent stem cells.

**Highlights:**

- Brain:satellite cell clock communication restores rhythms of core clock machinery in quiescent satellite cells
- Brain inputs are the dominant regulator of transcript rhythms in SCs, driving the oscillation of lipid metabolic genes.
- Autophagy in satellite cells is required for the oscillation of lipid metabolic genes.
- Early phases of muscle regeneration depend on brain-driven circadian signals.

## Introduction

The Earth’s 24-hour rotation generates a light-dark cycle to which most organisms’ behavior is synchronized. This synchronization is mediated by circadian clocks which allow anticipation of cyclic environmental changes^1^. In mammals, the central clock is found in the hypothalamic suprachiasmatic nucleus (SCN), it receives light inputs from the retina and subsequently sends entrainment signals to regularly reset clocks in peripheral tissues^2^. Recent evidence suggests that most peripheral clocks can oscillate, to varying extents, without inputs from the central SCN clock^3–10^. In these conditions, however, peripheral clocks have a limited ability to drive transcriptional output, suggesting that SCN clock outputs synergize with peripheral clocks to support a more complete circadian function in each tissue^5–8^. Particularly for peripheral tissues, changes in the external environment (such as food availability or diet composition) can alter circadian output^11–13^ as can signals emanating from other peripheral clocks^5,14–16^.

We have recently demonstrated the importance of circadian communication between the brain and the whole muscle for its normal muscle function and homeostasis^6,7^. Because muscle tissue is routinely damaged through daily activity, homeostasis is also dependent on satellite cells (SC), the resident stem cells of muscle which, in the adult, remain quiescent unless activated to repair damage. Previous work has shown that quiescent SCs have circadian transcriptional rhythms and exhibit day-night differences in autophagy, a macromolecule recycling process required for SC homeostasis and activation^17–21^. This cycling effect is lost during aging,^17^ during which time there is an extensive rewiring of the circadian transcriptome and a general decline in autophagy^18,22,23^. Indeed, autophagy-deficient SCs accumulate ROS, damaged mitochondria, and lose regenerative capacity^20^. While various whole tissues, including skeletal muscle^6,7,24,25^, receive key circadian signals from the central clock, it is less clear how individual cells, particularly stem cells, interact with peripheral and central clock inputs. Here, we investigated the communication between central and local SC clocks in SC circadian function. We demonstrate that reconstitution of the SC local clock alone has minimal impact on circadian SC regulation and homeostasis, while central clock inputs (even in the absence of the local clock) drive the oscillation of metabolic genes and sustain the early phases of muscle regeneration. Further, we demonstrate that metabolic gene oscillation is lost in SCs lacking autophagy, indicating that autophagic control in SCs is a key downstream mediator of central clock signals.

## Results

### Brain:satellite cell clock communication restores the rhythmicity of the core clock machinery

To investigate the contribution of local and central inputs on SCs’ circadian function, we re-expressed the essential clock gene *Bmal1* (also known as *Arntl*) specifically in SCs in mice that are otherwise *Bmal1* deficient (*Bmal1-*Stop^FLFL^; *Pax7*CreER^+/-^) (SCRE)^3,4^. We used WT littermates and complete *Bmal1*-KO mice (*Bmal1-*Stop^FLFL^) (KO) as positive and negative controls. In addition, we generated brain-specific reconstituted (*Bmal1*-Stop^FLFL^; *Syt10*^+/-^) (BRE)^26^, and brain plus SC-specific reconstituted (*Bmal1*-Stop^FLFL^; *Pax7*CreER^+/-^; *Syt*10^+/-^) (B+SC-RE) mice (Fig 1A). 6-week-old mice were injected with tamoxifen for 4 consecutive days (Fig S1A) to re-express *Bmal1* by inducing *Pax7*CreER activation and excision of the stop cassette within the *Bmal1* in SCs. All mice were housed in a 12:12 hour (h) light:dark cycle with ad libitum access to food and water. To avoid the potential influence of the severe muscle phenotypes of global *Bmal1*-KO^27^, all mice were sacrificed at 11-12 weeks of age. SCs from skeletal muscles were isolated by fluorescence-activated cell sorting (FACS) upon staining with specific antibodies for the detection of quiescent SCs from each genotype (Fig S1B), and restoration of BMAL1 protein was confirmed in SCs from SCRE and B+SC-RE mice as expected (Fig S1C). The absence or presence of *Bmal1* did not affect SC number (Fig S1D).

**Figure 1.**
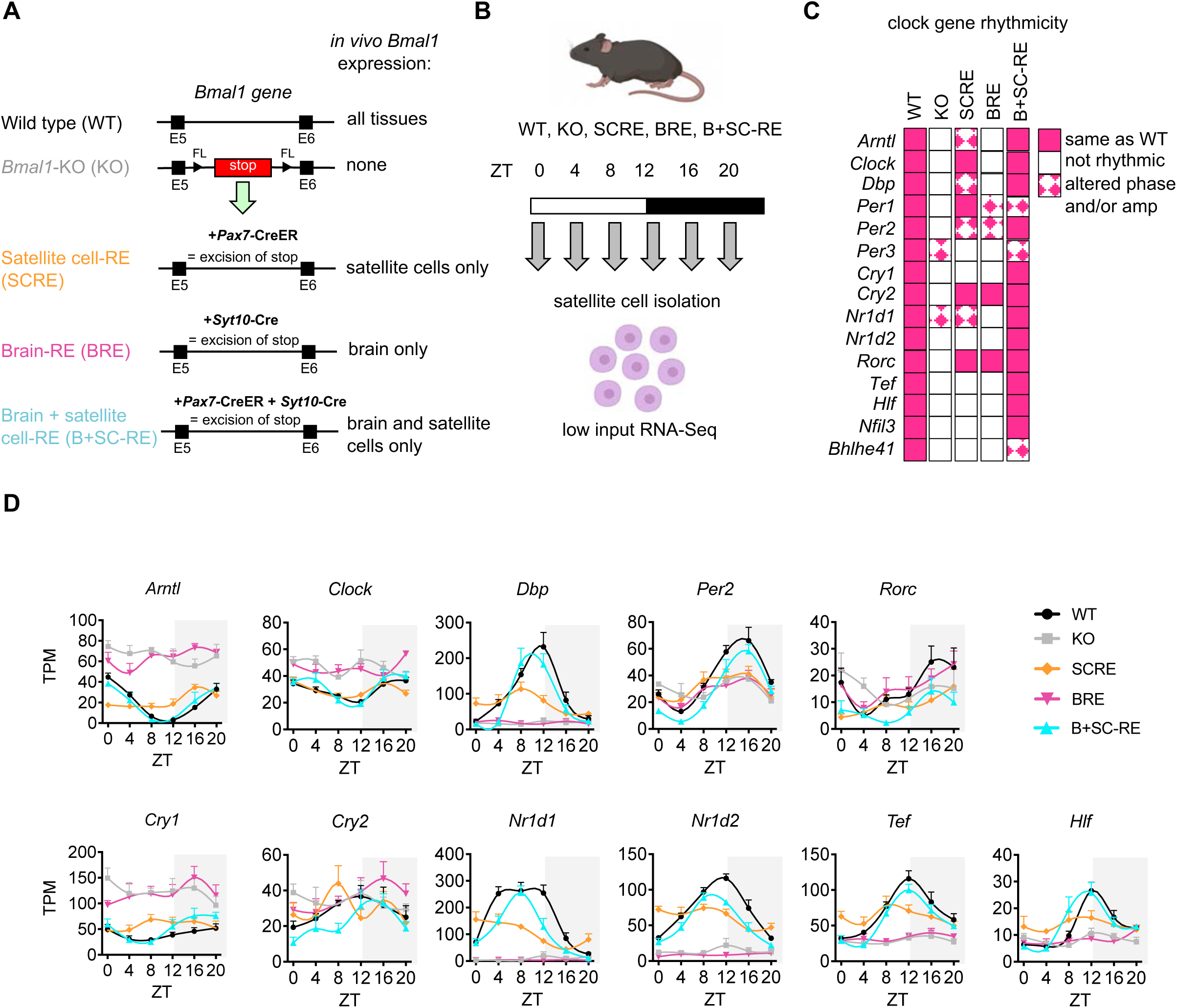
Dual restoration of *Bmal1* in brain and satellite cells is required for satellite cell clock rhythmicity. **A**. Mouse genetic scheme. **B**. Experimental workflow. Light and dark bars refer to 12-hour light and 12-hour dark phase, respectively. n of 6 mice (3 males and 3 females) per timepoint per genotype (also applies to C-D). **C**. DryR rhythmicity analysis of core clock genes in satellite cells. Only genes designated as rhythmic in WT satellite cells were considered. **D**. Clock gene expression in satellite cells. *Arntl* is also known as *Bmal1; Nr1d1* and *Nr1d2* are also known as *Reverbα* and *Reverbβ* respectively. TPM-Transcripts Per Million mapped reads, ZT-Zeitgeber Time. Figure 1B was created using BioRender.com. See also Figure S1. Wild-type (WT), Bmal1 total knock-out (KO), Bmal1 reconstitution in satellite cell only (SCRE), Bmal1 reconstitution in brain only (BRE), Bmal1 reconstitution in brain and satellite cells (B+SC-RE).

To assess the 24 h SC transcriptome of each genotype, we FACS-isolated SCs (Fig S1B) every 4 h, for a total of 6 time-points (Fig 1B) and performed low-input RNA sequencing (RNA-seq)^28^. We then assessed clock function by analysis of core clock gene oscillations in the 5 genotypes (Fig 1C,D and S1E). As expected, the clock was severely disrupted in SCs from KO mice. SC-specific restoration of *Bmal1* (SCRE) resulted in 8 out of 15 core clock genes oscillating but only four had the correct phase and amplitude (*Clock, Per1, Cry2, Rorc*). Hence, the local SC clock has limited autonomy under these conditions. When *Bmal1* was solely restored in the brain (BRE), only 4 out of 15 core clock components oscillated in SCs (*Per1, Per2, Cry2* and *Rorc*) and only *Cry2* and *Rorc* exhibited the correct phase and amplitude. Interestingly, when *Bmal1* was simultaneously restored in the brain and SCs (B+SC-RE), the majority of core clock genes oscillated identically to WT (Fig 1C,D), except for three genes *Per1*, *Per3* and *Bhlhe41* (Fig S1E). These results suggest that local SC *Bmal1* is not sufficient to support normal clock oscillations in these cells, while the integration of central inputs (from BRE) by local SC *Bmal1* is necessary and sufficient to drive core clock rhythmicity (Fig 1C,D).

### Central clock-driven inputs restore metabolic gene oscillations in satellite cells independently of the local clock

To identify the contributions of the local SC clock and central clock, and their interdependency, in the regulation of daily transcriptional rhythms, we compared the circadian transcriptome of SCs from the five genotypes using the circadian algorithm DryR^29^ (Fig 2A and S2, Table S1). Of the 960 oscillatory genes identified in WT SCs, 840 (87.5%) lose oscillation in *Bmal1*-KO mice. Restoring *Bmal1* solely in SCs rescued oscillation of only 326 (34%) genes, whereas restoring *Bmal1* in the brain rescued 572 genes (59.6%). Simultaneous reconstitution of *Bmal1* in the brain and SCs was not additive, with 564 genes (58.8%) rescued in this condition (Fig 2B). We next asked whether the SC clock may perform a gating function, as recently reported in whole muscle^7^, in which the local clock prevents *de novo* oscillations (not present in WT mice). However, *de novo* gene oscillations showing similar peak phase in SCs were detected to similar extents in BRE and B+SC-RE mice (Fig 2B and Fig S2A,B). To provide additional evidence, we also ran the JTK algorithm^30^ (Table S2) by using a low and a high statistical threshold and found a similar number of oscillating genes (Fig S3A). Focusing on genes that oscillate in WT and the respective genotype, we ranked DryR models based on the number of rescued oscillating genes in the same phase as in WT (Fig 2C). We identified 220 gene oscillations in the BRE and B+SC-RE groups, which we refer to as centrally driven since they oscillate in the BRE and do not require the local SC clock. Enrichment analysis by using the molecular signature database (MsigDB) identified that these centrally driven genes are enriched for terms involved in lipid metabolism (including sterol biosynthetic process and cholesterol biosynthesis) (Fig 3A). Similar results were found using DAVID gene ontology analysis (Fig S4A). We further tested whether the local clock is dispensable for oscillation of this gene set by using an inducible mouse model of *Bmal1* depletion in SCs (Bmal1^τιPax7ER^). DryR analysis of the SC circadian transcriptome from these mice revealed that lipid metabolic gene oscillations are independent of the SC clock (Fig 3B and S4B), providing independent evidence that these genes are not under local clock control. Indeed, lipid metabolism genes (*Apoe, Ldlr, Cyp51 and Fdft1*) follow very similar patterns in the two independent datasets (Fig 3C and S4C). Homer analysis predicted several transcription factors (TF) motifs that may be involved in mediating central clock-driven signals, including nuclear factor Y (NFY) which appears in both datasets (Fig 3D). Together, these data suggest that metabolic gene transcriptional oscillations are dependent on central inputs and independent of the local clock. This is in line with previous studies showing that the central clock is sufficient to drive rhythmic behavior such as feeding-fasting rhythms^26^, which drive oscillation of many genes in peripheral tissues even in the absence of the local clock^5,11,26,29^.

**Figure 2.**
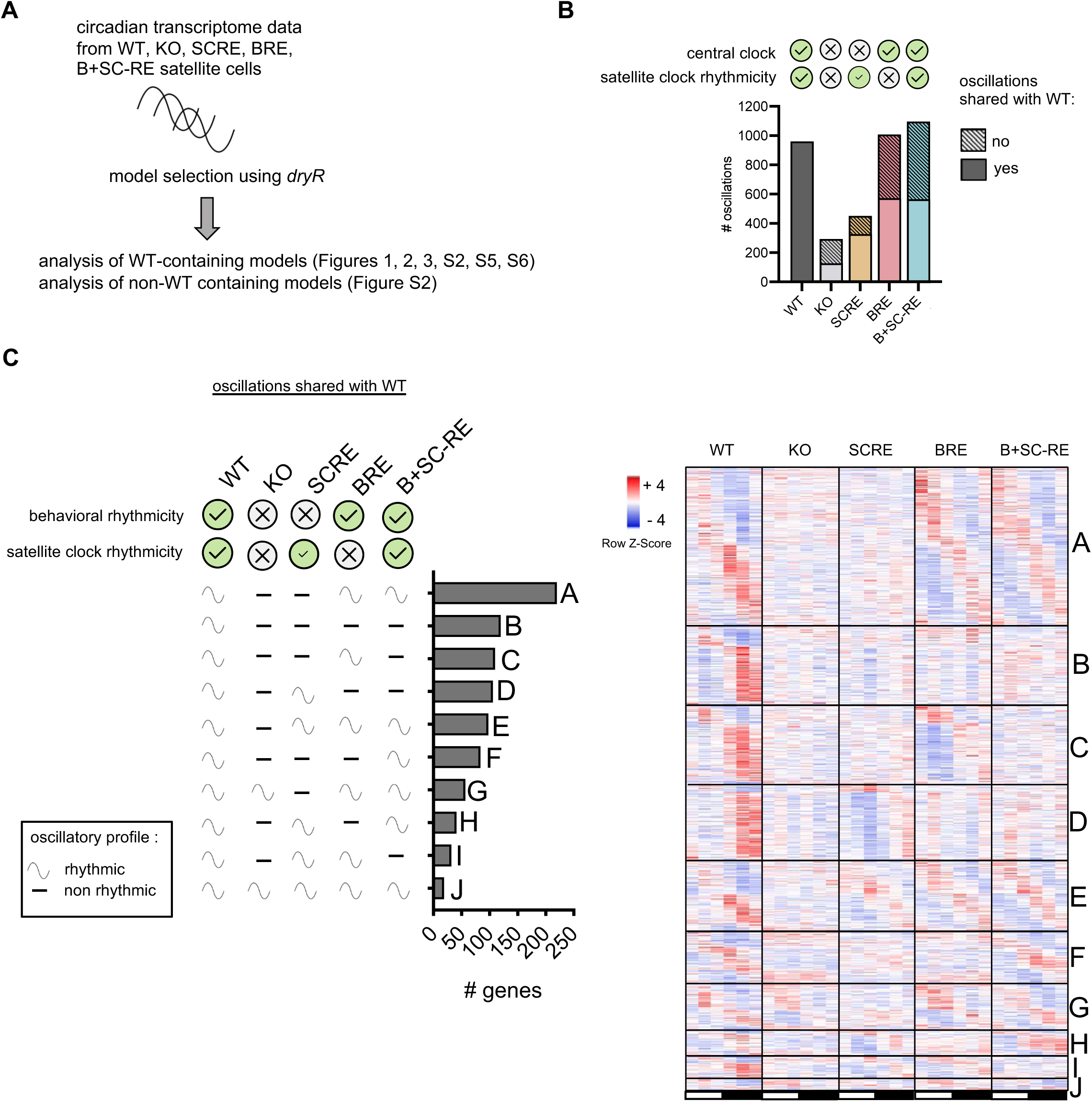
Determination of local and distal clock dependencies for transcriptional rhythms in satellite cells. **A**. Bioinformatic approach using dryR **B**. Rhythmic genes in common or not with WT (dryR BICW> 0.4, amp> 0.25, Cooks <1). Ticks and crosses refer to presence or absence of rhythmicity, as denoted. Small tick in SCRE relates to partial rhythmicity of core clock (see also Figure 1C) **C**. WT containing dryR models, and phase-aligned heatmap of mean expression of genes in each model over circadian time. White bar indicates light phase timepoints (ZT0, ZT4, ZT8) and black bar indicates dark phase timepoints (ZT12, ZT16, ZT20), ZT-Zeitgeber Time. Wild-type (WT), Bmal1 total knock-out (KO), Bmal1 reconstitution in satellite cell only (SCRE), Bmal1 reconstitution in brain only (BRE), Bmal1 reconstitution in brain and satellite cells (B+SC-RE).

**Figure 3.**
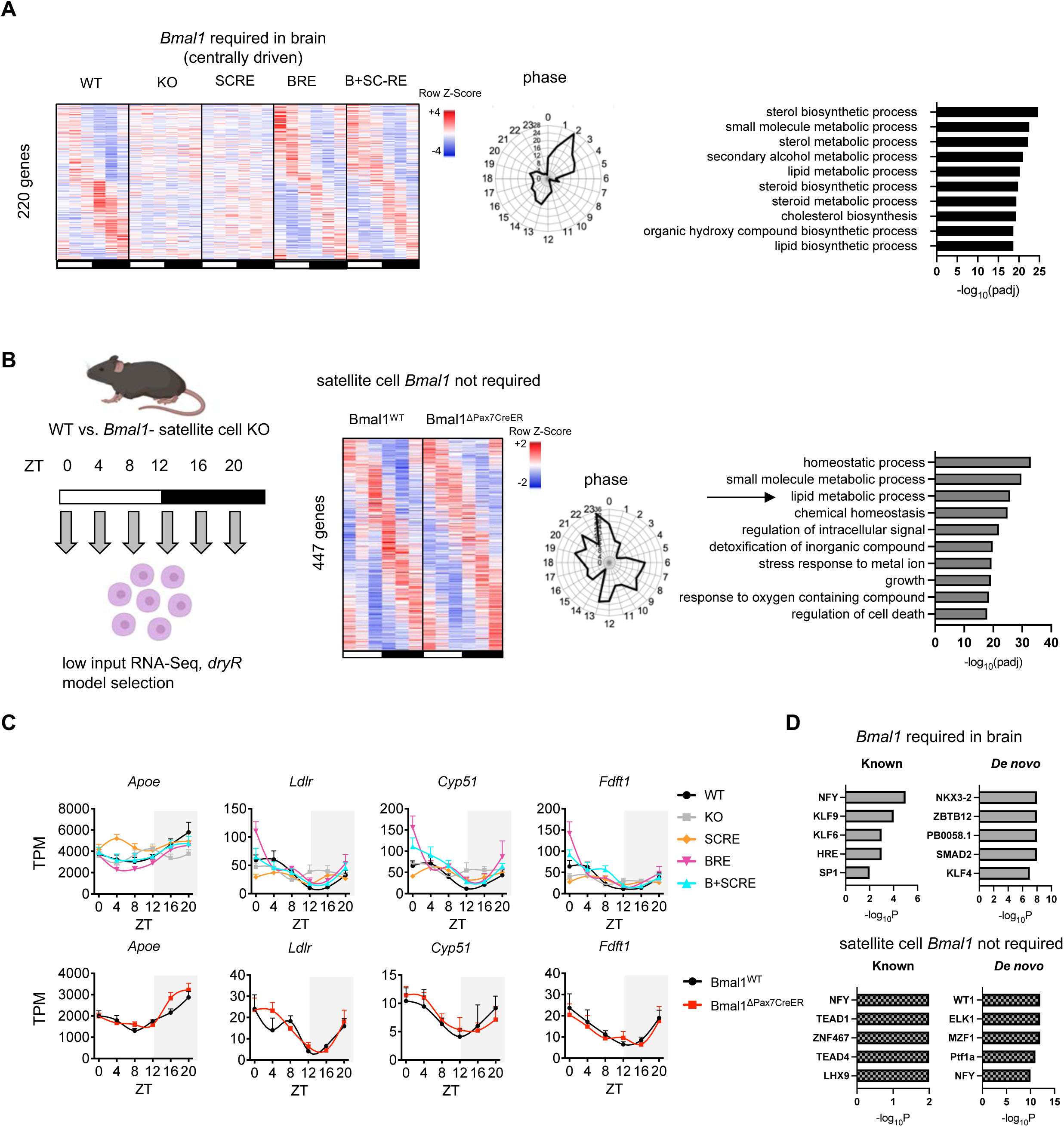
Central clock-driven signals are dominant for 24h-transcriptional rhythms of lipid metabolic genes in satellite cells. **A**. Heatmap and phase-plot from dryR model representing common oscillating genes in WT, BRE and B+SC-RE defined as centrally driven clock (BICW>0.4, Amp>0.25,Cooks<1) and MSigDB pathway enrichment for centrally driven genes. Numbers on the perimeter of phase plots refer to ZT, numbers on internal grid to the number of oscillatory transcripts (also applies to phase plot in B). **B**. Circadian experiment scheme, heatmap, phase-plot and pathway enrichment from dryR model of common oscillating genes in Bmal1^WT^ and Bmal1^1Pax7CreER^ satellite cells (BICW>0.6, Amp>0.25,Cooks<1) and MSigDB pathway enrichment. **C**. Examples of oscillating genes belonging to pathways found in the enrichment analysis in **A** and **B**. **D**. Transcription factor analysis of the 2 datasets by Homer. TPM-Transcripts Per Million mapped reads, ZT-Zeitgeber Time. Wild-type (WT), Bmal1 total knock-out (KO), Bmal1 reconstitution in satellite cell only (SCRE), Bmal1 reconstitution in brain only (BRE), Bmal1 reconstitution in brain and satellite cells (B+SC-RE).

### Autonomous and peripheral clock inputs influence SC transcriptional rhythmicity

In addition to the centrally-driven signals, we also tested a potential ‘autonomous’ role of local *Bmal1* to support transcriptional oscillations in SCs. We identified 41 output genes that oscillate in WT, SCRE, and B+SC-RE (Fig S5A). These output genes are involved in pathways associated with cell adhesion and cell projection (Fig S5A,B) and TF prediction analysis detected RFX5, ZBTB7A, TEAD4, GRHL2 and SMAD4 as potential regulators (Fig S5C). In specific dryR models (Table S3), only 84 genes were identified that oscillate specifically in WT and B+SC-RE (Fig 2C and S6), which suggests a limited role of SC core clock gene rhythmicity (Fig 1C,D) in driving circadian transcriptional output. These 84 genes, that require an ‘integration’ of *Bmal1* signals from the brain and SC cells, are involved in tissue development and rhythmic process (Fig S6A,B) and are predicted to be regulated by TFs including MYF5, MYOG, TEAD1, AP4, and USF2 (Fig S6C). Rather than driving rhythms or integrating central clock signals, we reasoned that the SC clock may instead be required to integrate inputs coming from other peripheral, non-brain clocks. In support of this, 238 out of 685 (34.7%) oscillating genes are lost in SCs upon local *Bmal1* deletion in otherwise WT mice (Bmal1^τιPax7ER^) (Fig S5D). Hence, *Bmal1* in SCs is necessary for their oscillation. The local clock-dependent genes (lost in Bmal1^τιPax7ER^) were identified as involved in the regulation of signaling (Fig S5D) and TF binding site predictions identified P73, PAX8, SMAD3, E2F3, EGR2, and NRF2 as potential regulators (Fig S5E). Combining this with the limited role of the SC clock to drive rhythms of output genes in B+SC-RE, this argues that the clock is required to integrate peripheral signals for full rhythmicity in SCs.

### Autophagy is required for metabolic gene oscillations

Autophagic activity varies over the day^17^ and preliminary evidence suggests autophagy and circadian rhythms regulate one another^31–34^. In support of this, we found that macroautophagy regulation was overrepresented in centrally driven genes (Fig S7A). To directly test the role of autophagy on gene oscillations, we used an inducible model to deplete the autophagy regulator Atg7 specifically in SCs (Atg7^τιPax7ER^)^20^ and performed RNA-seq on Atg7^WT^ and Atg7^τιPax7ER^ SCs harvested over circadian time every 4 h. DryR (Table S4) analysis revealed that 139 out of 1044 (13.3%) oscillating output genes depended on autophagy to maintain rhythmicity (Fig 4A, Fig S8) and a subset of these autophagy-dependent genes are involved in lipid and cholesterol metabolism (Fig 4B). Metabolic genes with oscillations under autophagic control include *Apoe*, *Pdk1*, *Psmd8*, *Gdpd3,* and *H6pd* (Fig 4C). Notably, we also identified *Apoe* as a central clock-driven gene oscillating independently of the local clock (Fig 3C). In SC lacking autophagy, the core clock genes were still operational, 8 out of 11 genes showing similar oscillations as in WT, yet *Cry2* oscillation was disturbed, along with amplitude changes in *Dbp* and *Hlf* (Fig S7B). Additionally, *Rorc* had increased amplitude and was phase-shifted compared to WT. Further pointing to an interplay between autophagy and circadian output, 386 genes that do not normally oscillate acquired a *de novo* circadian pattern in SCs from Atg7^τιPax7ER^ mice underlying a possible gating function of autophagy on the circadian output (Fig S8B).

**Figure 4.**
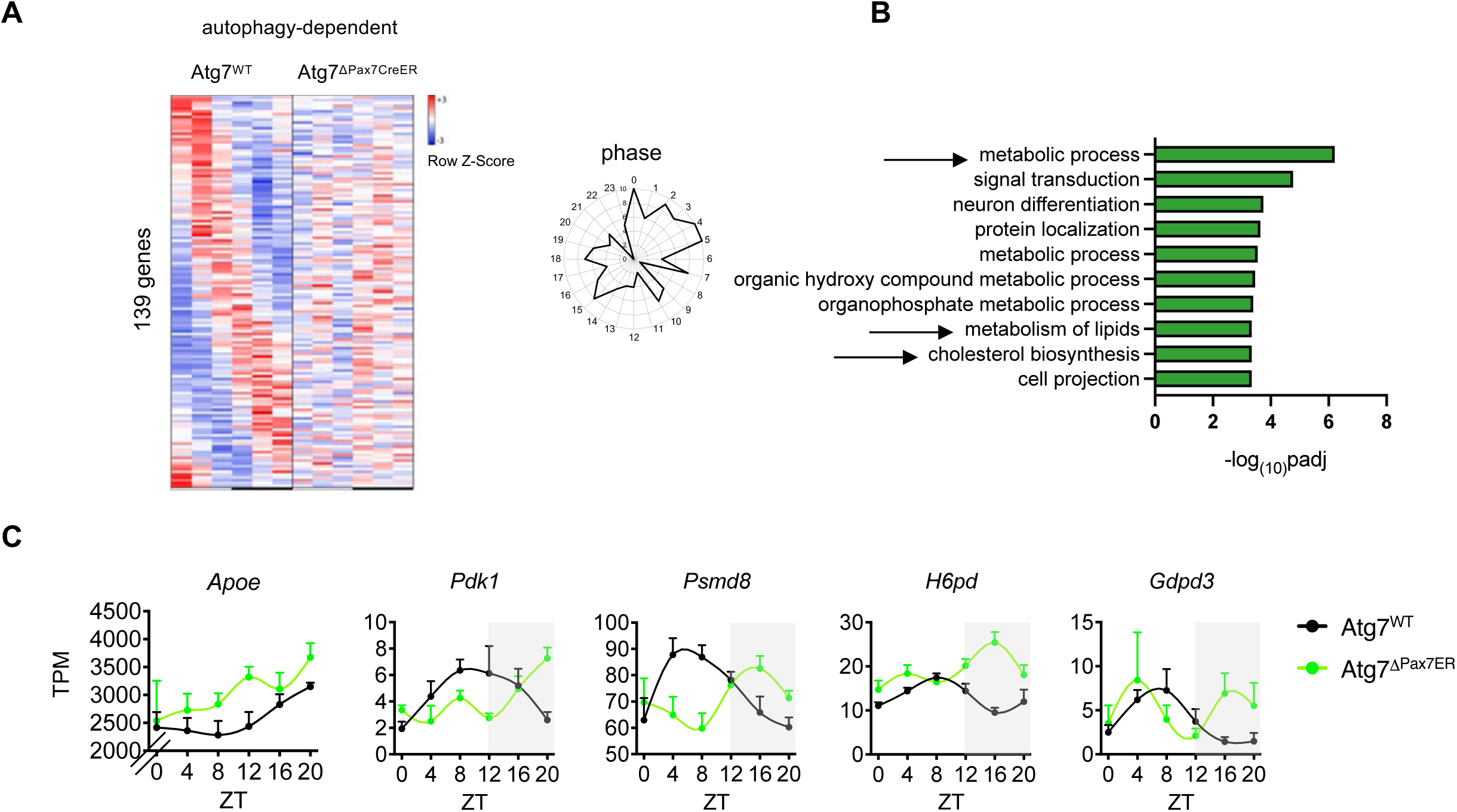
Autophagy controls oscillations of metabolic genes. **A**. Heatmap and phase-plot from dryR model showing 139 oscillating genes found only in Atg7^WT^ satellite cells (BICW>0.6, Amp>0.25,Cooks<1). Numbers on the perimeter of phase plots refer to ZT, numbers on internal grid to the number of oscillatory transcripts. **B**. Pathway enrichment of autophagy-dependent genes using MSigDB **C**. Examples of oscillating genes belonging to the category metabolic process from the pathway enrichment analysis. TPM-Transcripts Per Million mapped reads. ZT-Zeitgeber Time.

### Central clock signals support satellite cell metabolism and the early phases of muscle regeneration

To investigate the SC metabolic state, we used a recently developed low-input metabolomics protocol, ideal for rare cell types^35^ in SCs from WT and *Bmal1*-KO mice. Measured at the start of the light-dark cycle, we found an increased abundance of amino acids in SCs from KO mice compared to WT, and decreased carnitines, which are involved in lipid transport into mitochondria (Fig 5A,B). These results indicate a shift from lipid metabolism to protein degradation in SCs, which may be due to a decreased capacity to shuttle lipids into mitochondria for catabolism. To probe the functional relevance of this metabolic shift, we challenged SC function by cardiotoxin injury in the tibialis anterior (TA) muscles of the five genotypes. Cross-sectional area (CSA) of newly regenerated myofibers was performed four days post-injury (4DPI), a time point in which SCs are heavily involved in muscle repair. As expected^36,37^, we observed reduced CSA in *Bmal1*-KO mice versus WT (Fig 5C), an effect that was rescued in BRE, but not SCRE mice. Likewise, the frequency distribution of CSA showed a shift towards smaller fibers in KO and SC-RE mice (Fig 5D,E). Overall, these results further demonstrate the importance of central clock-driven inputs on SC physiology.

**Figure 5.**
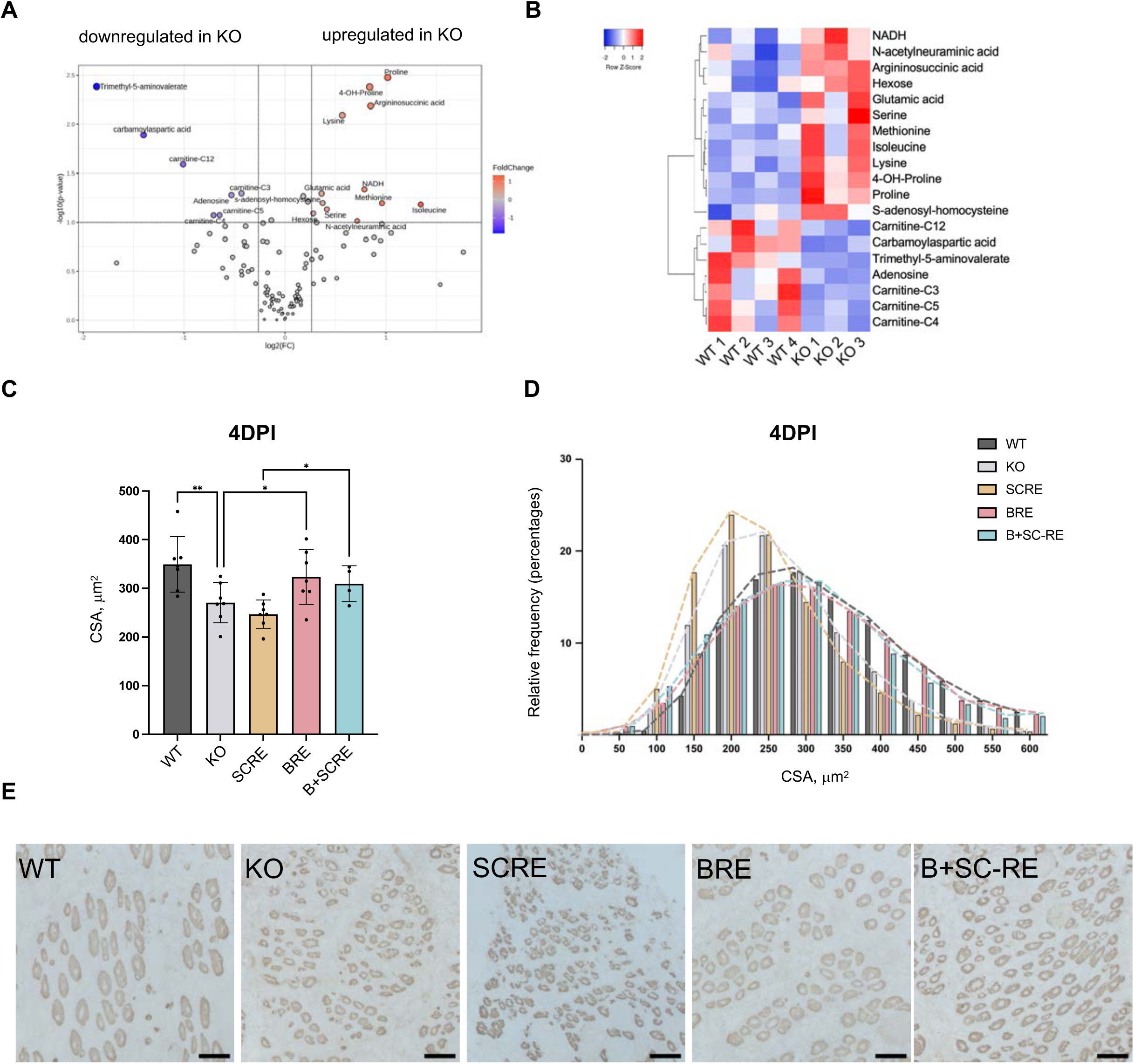
Disturbed metabolic homeostasis in KO SCs and rescue of regenerative defects by central-clock dependent signals. **A**. Volcano plot of metabolites detected in SCs from Bmal1 - WT and KO mice collected at ZT0. **B**. Heatmap of significantly affected metabolites in SCs from WT and KO mice collected at ZT0. **C**. Quantification of cross-sectional area (CSA, μm^2^) and **D**. Frequency distribution (%) from OCT sections of TA 4DPI, mice were injured and euthanized at ZT4. **E**. Representative images from eMHC staining in OCT sections of TA 4DPI from WT, KO, SCRE, BRE and B+SC-RE mice. Scale bar = 50µm. Resulted are displayed as means ± SEM P values are from one-way ANOVA uncorrected Fisher’s LSD *P < 0.05; **P < 0.01 ZT-Zeitgeber Time, tibialis anterior (TA), day post injury (DPI), embryonic myosin heavy chain (eMHC), wild-type (WT), Bmal1 total knock-out (KO), Bmal1 reconstitution in satellite cell only (SCRE), Bmal1 reconstitution in brain only (BRE), Bmal1 reconstitution in brain and satellite cells (B+SC-RE)

## Discussion

By specifically restoring BMAL1 in brain and satellite cells in mice otherwise lacking BMAL1, we determined how the central clock and non-dividing (quiescent) stem cells communicate circadian information for the first time. We found that limiting the reconstitution of *Bmal1* to SCs could not restore all core clock gene rhythms, and the few that did oscillate had abnormal amplitudes. This finding contrasts with epidermal cells, where local epidermal *Bmal1* expression has been shown to support most core clock gene rhythms and a large subset of transcriptional output genes, without central input^4,8^. SCs however, require local and brain *Bmal1* expression for core clock gene oscillations, suggesting a stronger dependence on central signals for normal circadian function in SCs than other cell types. Interestingly, however, restoring SC clock gene oscillations (by reconstituting *Bmal1* in the brain and SC) did not drive additional rhythmic gene output of non-core clock genes. Our finding that nearly half (44%) of transcriptional oscillations are lost in SCs from SC-specific *Bmal1* KO mice suggests that whilst *Bmal1* is not sufficient for the full repertoire, it is necessary for the oscillation of a substantial fraction of genes. We speculate that the quiescent SC clock may integrate signals from other peripheral clocks, to allow the full set of physiological circadian oscillations in these cells. SCs reside in a dynamic specialized niche below the basal lamina, where they interact with various stromal cells, muscle fibers, and the extracellular matrix^38–41^. This niche contributes to SC homeostasis during quiescence and supports the process of muscle repair. It is, therefore, tempting to speculate that these surrounding niche cells could also contribute to the satellite cell circadian output. In agreement, several reports suggest that peripheral clocks participate in both inter-cellular^15,42,43^ and inter-tissue communication^5,6,14,16^, and argue in favor of a ‘federated’ clock system^44^.

Among the different reconstitutions, we found that centrally-driven inputs restored the largest subset of gene oscillations in SCs, which may be related to feeding–fasting daily rhythmic behavior^11,26^. This may also explain our observation that centrally driven genes in SCs are mainly involved in lipid metabolism and that carnitine levels were reduced in SCs from KO mice. Amino acid levels increased in SCs from KO mice, suggesting an adaptive switch to protein degradation, likely to meet energy demands in the presence of altered lipid catabolism. These *Bmal1*-dependent changes in the quiescent state are similar to those reported in activated SCs from SC-specific *Bmal1*-KO^37^, in which decreased carnitines and increased amino acid dipeptides are detected. Our data suggests that *Bmal1* regulates these processes even before SC activation. We found that after muscle injury, muscle fibers with a restored central clock were more similar to WT than *Bmal1*-KO, or local *Bmal1-*SC-RE mice, which argues that central clock-driven signals can partially overcome the functional impact of this SC dysregulation.

Although our data show that local SC clock deletion did not affect the oscillation of central clock-driven metabolic genes, their oscillations were strongly inhibited in SCs lacking autophagy. Previous work has suggested autophagy may generate circadian transcriptional oscillations^31^ and autophagy was shown to have a direct role in mediating the beneficial effects of time-restricted feeding in flies^45^. Our results that autophagy-deficient SCs lose metabolic gene oscillations, even in the presence of central clock-driven inputs suggest autophagy is required to integrate the central inputs in this context. Our data further suggest that autophagy might also inhibit rhythmic gene expression, as a considerable number of typically non-rhythmic genes acquired oscillation in autophagy-deficient SCs. Considering SCs from aged mice exhibit altered day-night autophagy rhythms along with reprogrammed circadian output, the decrease of baseline autophagy during aging^20,23^ may be connected to the rewiring of circadian output. Future work remains to define the molecular mediators that underlie this form of cellular autophagy and inter-tissue communication and explore whether this is a general mechanism for quiescent stem cells in other tissues.

### Limitations of the study

Here, we use *in vivo* deletion and Cre-dependent reconstitution of the core circadian gene *Bmal1* based on our previously validated system^3–8,26,46^. In this study, we used a tamoxifen-inducible Cre to avoid restoring *Bmal1* in satellite cells during developmental stages, which may have resulted in nascent *Bmal1*-positive muscle fibers over time and hence obscured the interpretation of the satellite cell results. Nevertheless, the phenotype present in the conventional *Bmal1*-KO mice^27^ could be attributed, in part, to the developmental effects of *Bmal1* before 3 months of age^47^. Dissecting the developmental and circadian roles of *Bmal1* remains a challenge of the circadian field. We took care to harvest samples before 13 weeks of age, prior to the development of muscle fibrosis phenotypes^27^. Additionally, *Bmal1* has reported ‘moonlighting’ functions^48^ separate from its circadian role which our current methodology is unable to stratify. The rescue of muscle fiber size following *Bmal1* restoration in the brain, in addition to SC-specific effects, could also be related at least in part to effects on other cells that play an important role during muscle repair, such as cells in the niche and immune cells such as neutrophils and macrophages that are recruited to the injury site in the early phase of muscle regeneration^49^.

## Materials and methods

### Animal models

All mice were on a C57/BL6 background and housed at the Barcelona Biomedical Research Park under 12 hour light: 12 hour dark conditions with ad libitum access to normal mouse chow and water. For the Bmal1 tissue-specific restoration experiments, we crossed Bmal1 conditional gene-trap mice^4^ (Bmal1-stopFL), with the respective cre lines as mentioned in results, and used WT littermates from the same colony as controls. As required, Cre activity in SC was induced by daily intraperitoneal injections of tamoxifen (Sigma; 10 mg ml−1 in corn oil) at a dose of 5mg/25g bodyweight for 4 days at 6 weeks of age. RE Bmal1 mice were used up to a maximum of 12 weeks, to minimize impacts of reported early aging phenotypes in conventional Bmal1 KO mice evident at this point onwards^27^. Experiments were approved by the local ethics committee of the PRBB and Catalan government. The circadian dataset of autophagy-deficient SCs is from male mice, while datasets from RE models and specific *Bmal1* deletion in SCs, are generated from males and females mice.

### Satellite cell isolation by FACS

Following sacrifice, satellite cells were obtained from skeletal muscles as previously described^28^. Briefly, skeletal muscles from fore and hind limbs and abdominal muscles were harvested in 20 mL of DMEM plus 1% Penicillin–streptomycin (P-S) on ice, minced by blade, and digested at 37 °C for 1 h in DMEM media containing 1% P-S, liberase (0.1 mg·g^-1^ muscle weight) (Roche, 5401020001; 5 mg·mL^-1^), 0.3% dispase (Gibco, #17105-041), 0.5 μM CaCl2, and 6 μM MgCl2. The reaction was stopped by addition of at minimum, an equal volume of cold DMEM, 1% P-S, and 10% FBS (Sigma-Aldrich F7524). Cells were filtered sequentially through a 100-μm and 70-μm cell strainer, then pelleted by 4 °C centrifugation at 670g for 10 minutes. Cell pellets were resuspended in RBC lysis buffer 1× (eBioscience, 00-4333-57) for 5 min on ice, then enzymatic reactions were stopped by with addition of PBS, and cells filtered through a 40μm cell strainer. After 4 °C centrifugation at 670g for 10 minutes, pellets were resuspended in 2.5% FBS in PBS. For FACS analysis, the following antibodies were used: PE-Cy7-conjugated anti-CD45 (Biolegend, 103114), anti-Sca-1 (Biolegend, 108114), PE-Cy7-conjugated anti-CD31 (Biolegend, 102418) were used for lineage-negative selection, and Alexa Fluor 647-conjugated anti-CD34 (BD Pharmigen, 560230), and PE-conjugated anti-α7-integrin (AbLab, 53-0010-05) were used for double-positive staining of quiescent satellite cells.

### Histology and immunohistochemistry (IHC) in muscle cryosections

TA muscles were snap-frozen or pre-fixed (2% PFA 2 h at 4 °C and washed in PBS), cryo-preserved and frozen in an isopentane/liquid nitrogen double bath, and stored at −80 °C until analysis. Immunochemistry and immunofluorescence staining were performed in 10 μm sections. Labeling of cryosections with embryonic myosin heavy chain was performed using the peroxidase M.O.M kit staining (Vector Laboratories, SK-4100) according to the manufacturer’s instructions. Double immunostaining was performed by sequential addition of each primary and secondary antibody using appropriate positive and negative controls. For snap-frozen samples, sections were air-dried washed on PBS, and permeabilized with ice-cold methanol before antigen retrieval. For antigen retrieval, the sections were incubated in boiling citric acid (0.01 M, pH=6) for 10 min, cooled, and washed with PBS. The samples were then incubated with primary antibodies for PAX7 and laminin according to the manufacturer’s instructions after blocking for 1 h at room temperature (RT) with 5% BSA and M.O.M blocking diluted in PBS (30 min). Subsequently, the slides were washed on PBS and incubated with appropriate secondary antibodies and labeling dyes. For immunofluorescence, secondary antibodies were coupled and nuclei were stained with DAPI (Invitrogen). After washing, tissue sections were mounted with Fluoromount G (SouthernBiotech).

### Low-input RNA sequencing preparation from FACS-sorted satellite cells and analysis

As outlined in Sica et al^11^, RNA from FACS-isolated SCs was extracted using an RNeasy micro kit (Qiagen), and quality control was performed using Agilent 2100 Bioanalyzer or fragment analyzer. RNA with RIN values of 7 or higher were used with a rRNA ratio (28S/18S) of 2 or above. 0.3 ng RNA was used for each sample to generate barcoded RNA-seq libraries using the NEBNext Single Cell/Low-Input RNA Library Prep Kit for Illumina (New England Biolabs). Library size was checked using Agilent 2100 Bioanalyzer and concentration determined Qubit® fluorometer (Life Technologies). Libraries were sequenced at 650 pM on a P3 flow cell of the NextSeq 2000 (Illumina) to generate 60-base single reads. bcl2fastq 2.20 Software (Illumina) was used to obtain FastQ files for each sample. We used Nextflow nf-core/rnaseq v.3.2 pipeline^31^ to map the raw reads to the reference genome using STAR ALIGNER^32^, project the alignments onto the transcriptome, and to perform the downstream transcript-level quantification with Salmon^33^. The R package TXIMPORT^34^ was used for gene-level summarization of read counts and transcript per million (TPM) abundances. Four to six replicates for each genotype and timepoint were used in each dataset.

### RNA-seq data processing

Raw FASTQ files were subjected to quality control (QC) procedures to trim low-quality bases, remove adapter sequences and filter out low-quality reads using FastQC v0.11.9 and BBDuk v35.85 with a minimum read length of 35 bp and a minimum quality score of 25. High-quality reads were mapped with STAR^32^ against the reference mouse genome (GRCh38). The number of reads mapped to genes was quantified using FeatureCounts and raw counts matrix was loaded into R (https://www.R-project.org/). to compute the Transcripts Per Million (TPM) values. Biological (sex) and technical (RNA extraction date) batch effects were corrected with the ComBat() function from sva R package.

### Circadian algorithmic analysis, pathway enrichment and transcription factor analysis

Differentially rhythmic categories corresponding to gain, loss or the same rhythm were defined using the “dryseq()” function of the R package dryR^29^, including technical and biological variables as potential batch effects. The following filtering criteria were used to select genes with the expression rhythmic in a respective model: Bayesian information criterion weight (BICW) ≥ 0.4, and amplitude ≥ 0.25. Genes with maximal Cook’s distance > 1 across the replicates were considered outliers and removed. Models with only one 1 gene were discarded. For DryR 2 group comparisons (Bmal1^WT^ vs Bmal1^τιPax7ER^ and Atg7WT vs Atg7 ^τιPax7ER^) BICW > 0.6, amplitude > 0.25 and Cook’s distance < 1 were used. For JTK_CYCLE p<0.05 or p<0.01 was used, as described in the figures. Enrichment analysis was performed using the Molecular Signatures Database^50,51^ (MSigDB R package) using hypergeometric tests (“phyper()” function from stats R package). False Discovery Rate (Benjamini-Hochberg) was computed. DAVID analysis^52,53^ was performed using the online tool and GOTERM_BP_DIRECT gene ontology was considered. For transcription factor (TF) predictions, the HOMER^54^ (Hypergeometric Optimization of Motif EnRichment) tool was used to find and annotate motifs for each DryR model independently. Statistically significant motifs were found by running ‘findMotifs.pl’ using the Ensembl^55^ Archive Release 104 reference as the background set of genes, focusing on a genomic region of 500 bps upstream to 100 bps downstream of the TSS to locate the motifs. *De novo* motifs with consensus sequences were annotated to specific genomic features (i.e., target genes) using ‘annotatePeaks.pl’.

### Targeted low-input metabolomics of polar metabolites

For low-input targeted metabolomics of polar metabolites, 5000 cells were sorted via FACS and processed according to previously published protocols with minor modifications^56,57^. Specifically, cells were sorted directly into 25 µl of extraction buffer that included 13C labeled yeast extract as an internal standard diluted in acetonitrile. The FACS sorting was performed using a 70µm nozzle with water containing 2g/L NaCl as sheath fluid. As a control, 5,000 events of cell debris were sorted. Additionally, a negative control only consisting of an equal amount of sheath fluid (5µl) was prepared. Metabolites were identified by their retention times and at least two known fragments. The height of chromatographic peaks was used as a measure of metabolite abundance.

### Processing of metabolomics data

For generation of the volcano plot, metabolite values (raw peak height) were processed using Metaboanalyst^58^ (metaboanalyst.ca, v6). Metabolites with missing values were excluded, values were log10 transformed and subject to auto scaling (mean-centered and divided by the square route of the SD of each variable). A raw P-value threshold of 0.1 a fold change (FC) threshold of 1.2 were used. Metabolites that passed this threshold were subsequently compared against recorded values from negative (no cell) controls, and metabolites detected <1.2 fold in experimental groups (either WT or KO) versus no cell controls were excluded (carnitine C6 and carnitine C8). For metabolites identified as regulated in KO vs WT SCs, lists of significant metabolites and original data were joined using Galaxy^59^ heatmaps were generated using Heatmapper^60^ using the following settings: clustering method; average linkage applied to rows, distance measurement method; Pearson).

### Western blotting

FACS-sorted satellite cells were lysed in RIPA buffer (50mM Tris HCL pH8, 150mM NaCl, 5mM EDTA, 15mM MgCl2, 1% NP40, 0.5% sodium deoxycholate in water with phosSTOP phosphatase inhibitor (Roche) and cOmplete Mini EDTA-free protease inhibitor (Sigma-Aldrich) added according to manufacturer’s instructions) on ice for 30 minutes. Samples were spun at 16,000g for 15 mins at 4°C, and supernatant saved. 5X Loading buffer (5% w/v SDS, 0.2M Tris, pH 6.8, 5% v/v Beta-mercaptoethanol, 0.15 mg/mL Bromophenol blue, 50% v/v glycerol) was added to a 1X concentration, and samples were incubated for 5 minutes at 95°C. Each sample was passed 3 X through 29G insulin syringe, then spun at 10,000g for 1 mins at RT and run on 4–20% Precast Protein Gels (Bio-Rad) with SDS-containing running buffer (0.1% SDS, 25mM Tizma base, 0.19M glycine in water). Samples were transferred to nitrocellulose membranes in transfer buffer (20% MeOH, 24.8 mM Trizma base, 191.8 mM Glycine in water) at 330mA for 1.5h. Membranes were incubated with ponceau (0.% w/v ponceau, 1% acetic acid in water) to visualise proteins, then blocked in 5% powdered milk in TBST (0.1% Tween, 10mM Tris, 100mM NaCL in water) for 1 hour. Blots were incubated overnight at 4°C with rabbit anti-BMAL1 (ab93806). After 30 minutes of TBST washes, blots were incubated with HRP-linked anti-rabbit secondary for 1 hour at RT, washed with TBST for 30 minutes, and imaged using SuperStrength West Femto ECL reagent and the BioRad ChemiDoc imaging system.

### Author contributions

V.S. and P.M.-C. conceptualized the work, designed the study, and supervised the overall experiments. V.S., E.A. and V.L. performed most experiments, V.S. J.G.S and O.D. performed the bioinformatics data analysis of the circadian transcriptomes, V.S. and J.G.S. interpreted the experimental data and wrote the manuscript with P.M.-C. and critical input from A.L.S and E.P. M.E. and N.C.-W. provided support for metabolomic experiments and analyses, S.A.B. provided the Bmal1-StopFL mice strain used in this study. Competing interest: A.L.S., E.P., and P.M.-C. are investigators of Altos Labs. S.A.B. is a cofounder and scientific advisor of ONA Therapeutics. The other authors declare no competing interests.

## Acknowledgments

We thank L. Ortet, J. Segalés and José M. Ballestero-Zambrano for their technical contributions. We are also indebted to A. Dopazo (CNIC-Genomics Facility) for help in transcriptomics, A. Rueda and A. Saera-Vila at Sequentia Biotech (Barcelona) for additional bioinformatic analysis, J. A. Fernández-Blanco (PRBB Animal Facility) for animal care. Funding: The authors acknowledge funding from MICINN-Spain (RTI2018-096068 to P.M.-C. and E.P.), ERC-2016-AdG-741966 to P.M.-C., ‘‘la Caixa’’ (HEALTH-HR17-00040). UPGRADE (H2020825825), and María-de-Maeztu Program for Units of Excellence to UPF (MDM-2014-0370) and Altos Labs Inc. Research in the J.G.S laboratory is supported by Agencia Estatal de Investigación (AEI) Aid RYC2022-035133-I funded by MICIU/AEI/10.13039/501100011033 and by the FSE+, and project PID2023-150233NA-100 funded by MICIU/AEI/10.13039/501100011033 and FEDER, EU, and AFM-Téléthon grant #28842. Research in the S.A.B. laboratory is supported by ERC-787041, the Foundation Lilliane Bettencourt, the Spanish Association for Cancer Research (AECC), and the Worldwide Cancer Research Foundation (WCRF). The IRB Barcelona is a Severo Ochoa Center of Excellence (SEV-2015-0505). V.S. acknowledges funding from FEBS long-term fellowship and European Union’s Horizon 2020 Research and Innovation Programme under the Marie Sklodowska-Curie grant agreement 895380.

---

**Figure S1.**
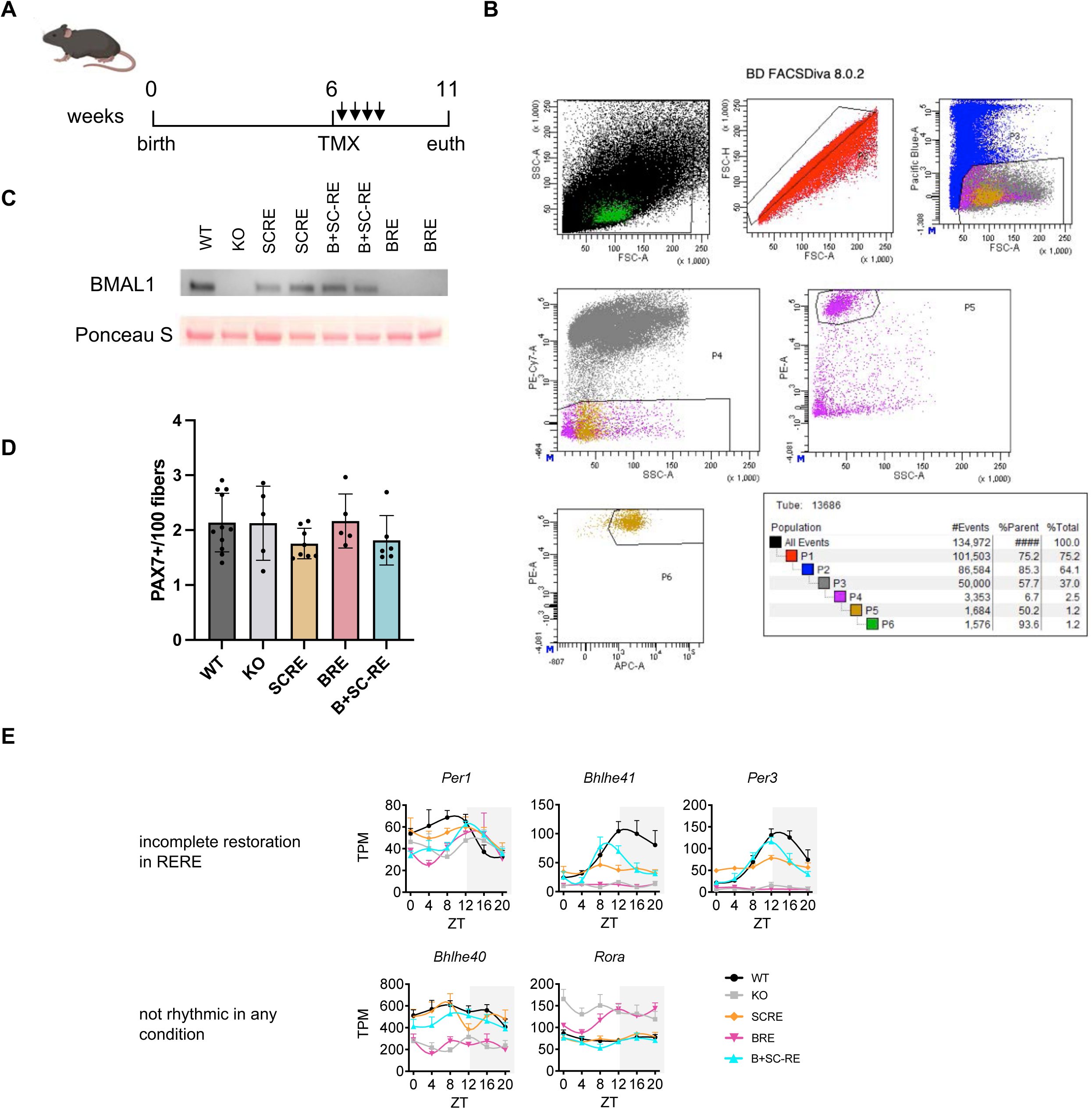
Characterization of satellite cell *Bmal1*-reconstituted models. **A**. Scheme of tamoxifen (TMX) administration for Pax7-CreER expression **B**. Representative image of satellite cell sorting strategy: PE-Cy7-conjugated anti-CD45, anti-Sca-1, PE-Cy7-conjugated anti-CD31 were used for lineage-negative selection, and Alexa Fluor 647-conjugated anti-CD34 (APC-A), and PE-conjugated anti-α7-integrin were used for double-positive staining of quiescent satellite cells. **C**. BMAL1 protein assessment of satellite cells from indicated genotypes **D**. Quantification of PAX7 immunofluorescence in OCT of tibialis anterior slice. **E** Clock genes not completely restored in B+SC-RE and non rhythmic in any condition. TPM-Transcripts Per Million mapped reads, ZT-Zeitgeber Time. Wild-type (WT), Bmal1 total knock-out (KO), Bmal1 reconstitution in satellite cell only (SCRE), Bmal1 reconstitution in brain only (BRE), Bmal1 reconstitution in brain and satellite cells (B+SC-RE).

**Figure S2.**
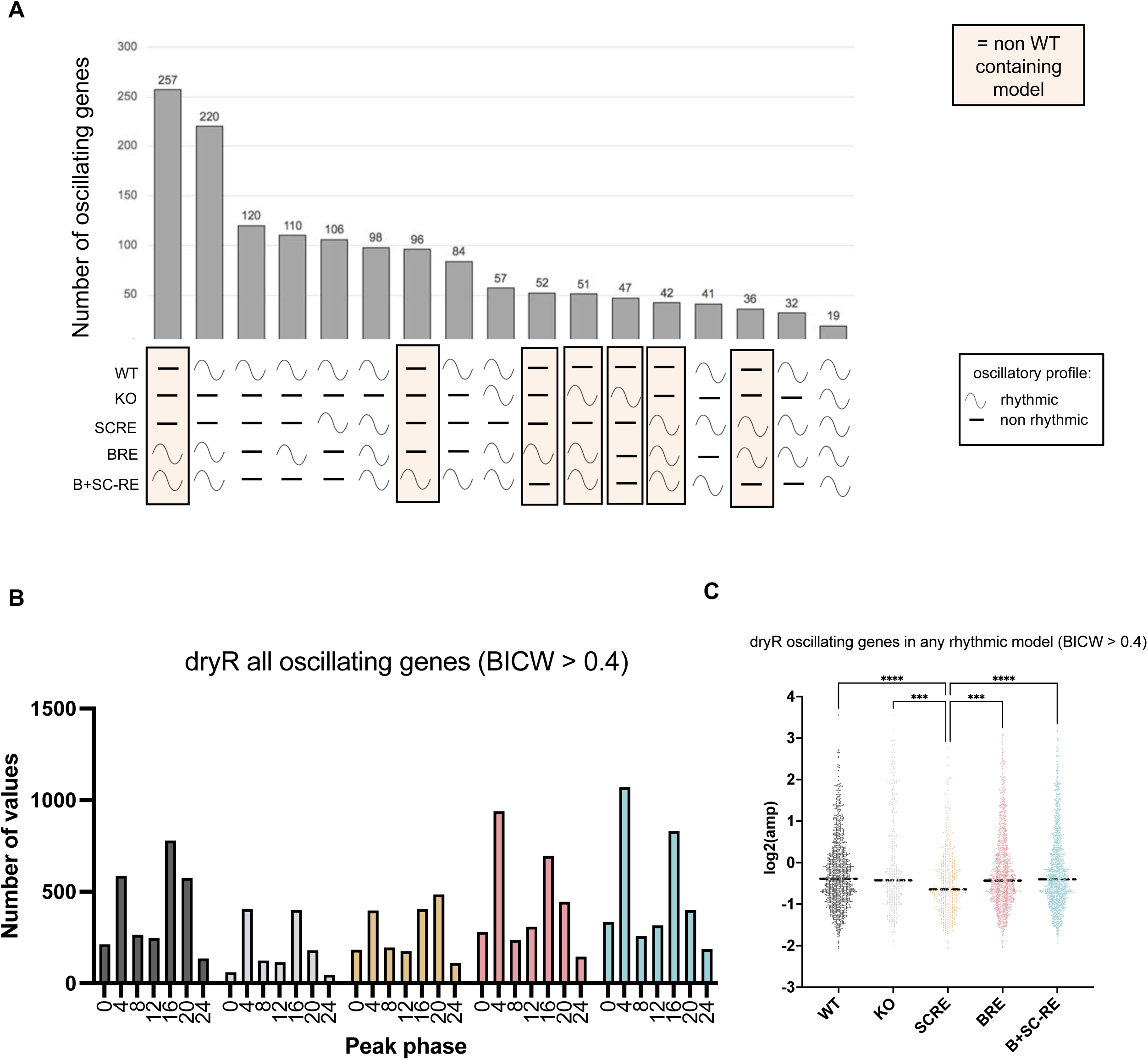
dryR bioinformatic analysis. **A**. Number of oscillating genes in each dryR models. Models in which WT is not included are designated with an orange box. **B**. Peak phase of genes classified as oscillating by dryR **C**. Amplitude analysis of genes classified as oscillating by dryR.

**Figure S3.**
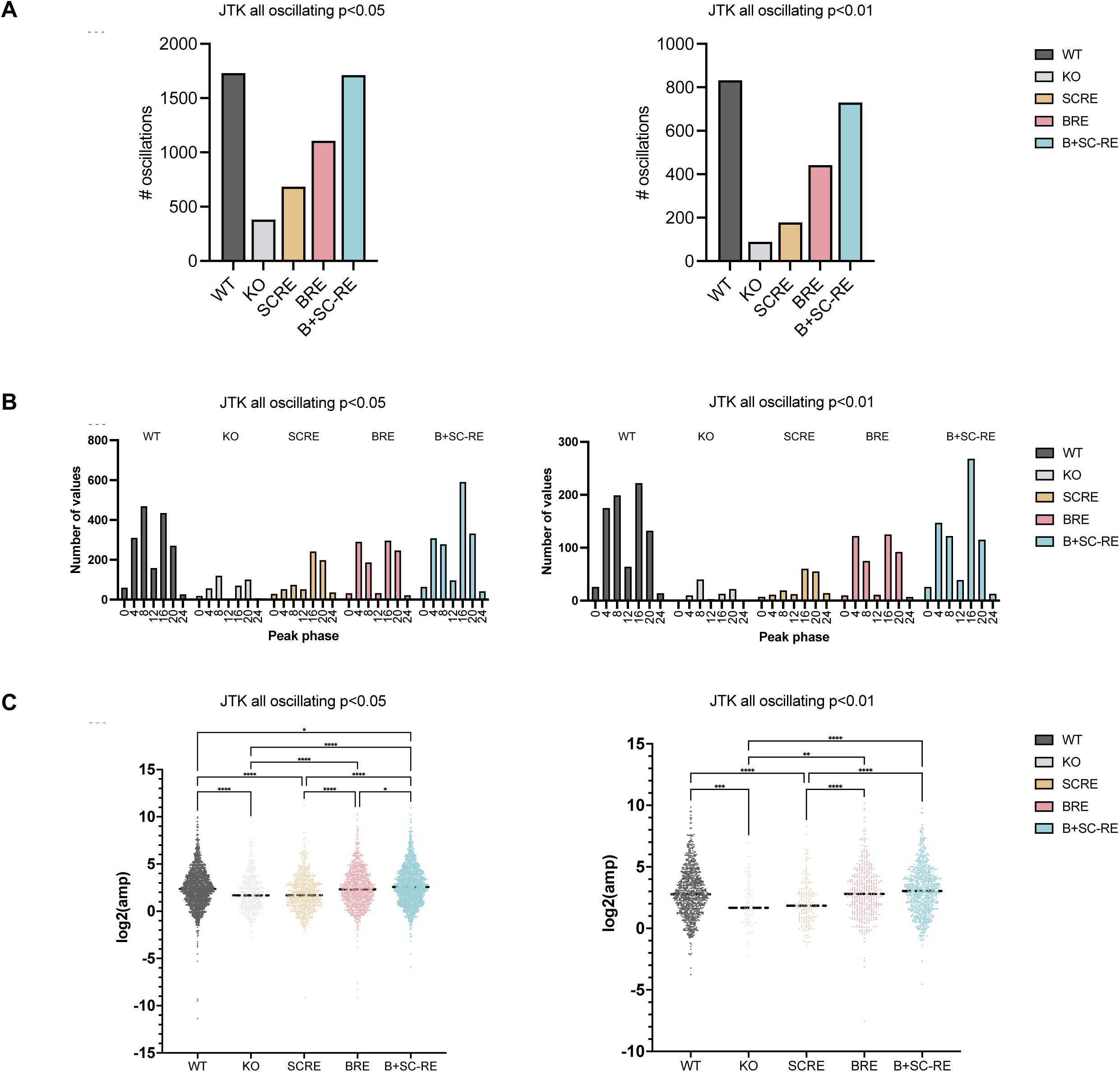
*JTK* analysis comparing low and high stringency strategies. **A** number of rhythmic genes in the indicated genotypes. **B**. Peak phase of genes classified as oscillating by *JTK.* **C**. Amplitude analysis of genes classified as oscillating by *JTK*.

**Figure S4.**
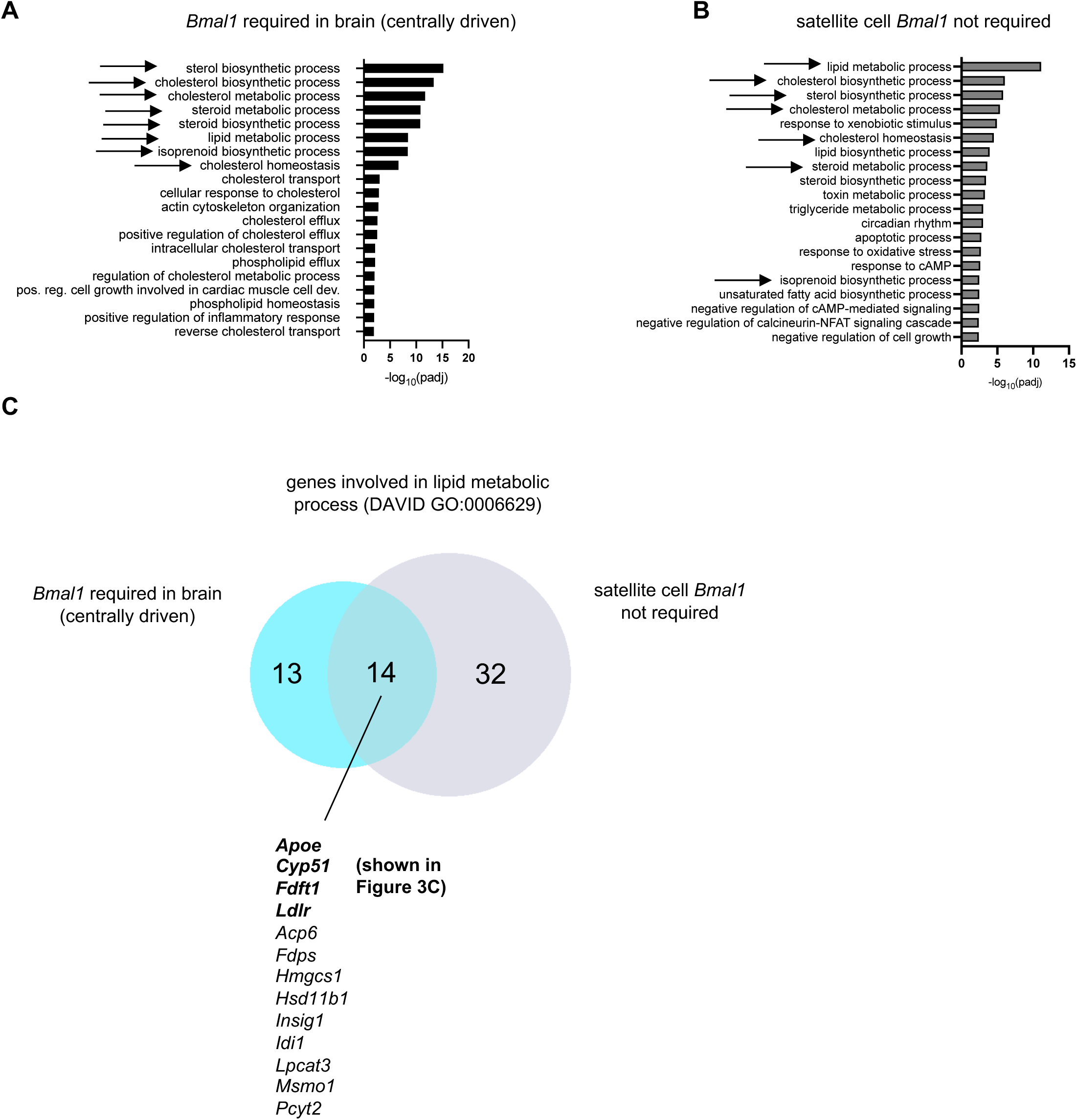
Centrally-driven genes in SC regulate lipid metabolism. **A**. DAVID gene ontology analysis for biological function for ‘*Bmal1* required in brain (centrally driven)’ gene set (see also Figure 3A). **B**. DAVID gene ontology analysis for biological function for ‘satellite cell *Bmal1* not required’ gene set (see also Figure 3B). For both A and B, arrows are terms present in both analyses. **C**. Venn diagram of lipid metabolic genes oscillating in ‘*Bmal1* required in brain (centrally driven)’ gene class and ‘satellite cell *Bmal1* not required’ DryR gene classes.

**Figure S5.**
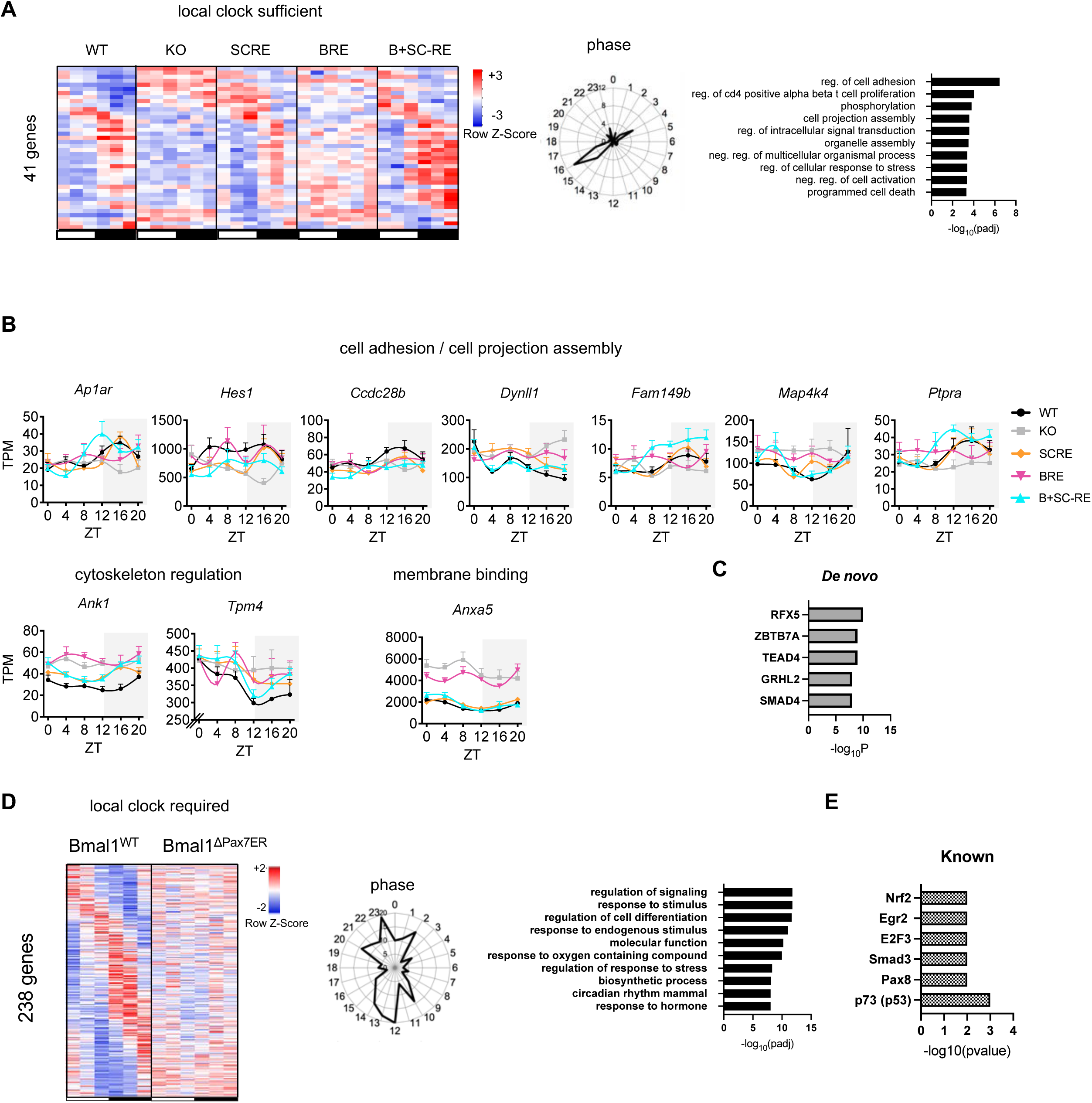
Satellite cell clock regulated oscillations. **A**. Heatmap, phase-plot and pathway enrichment (MsigDB) from dryR model representing common oscillating genes in WT, SCRE and B+SC-RE defined as autonomous clock (BICW>0.4, Amp>0.25,Cooks<1). Numbers on the perimeter of phase plots refer to ZT, numbers on internal grid to the number of oscillatory transcripts (also applies to phase plot in D). **B**. Oscillating genes belonging to pathways found in the enrichment analysis. **C**. Transcription factor analysis by Homer. **D**. Heatmap, phase-plot and pathway enrichment (MsigDB) from dryR model representing 238 gene oscillations that depend on the local clock (BICW>0.6, Amp>0.25,Cooks<1). **E.** Transcription factor analysis by Homer. TPM-Transcripts Per Million mapped reads, ZT-Zeitgeber Time. Wild-type (WT), Bmal1 total knock-out (KO), Bmal1 reconstitution in satellite cell only (SCRE), Bmal1 reconstitution in brain only (BRE), Bmal1 reconstitution in brain and satellite cells (B+SC-RE).

**Figure S6.**
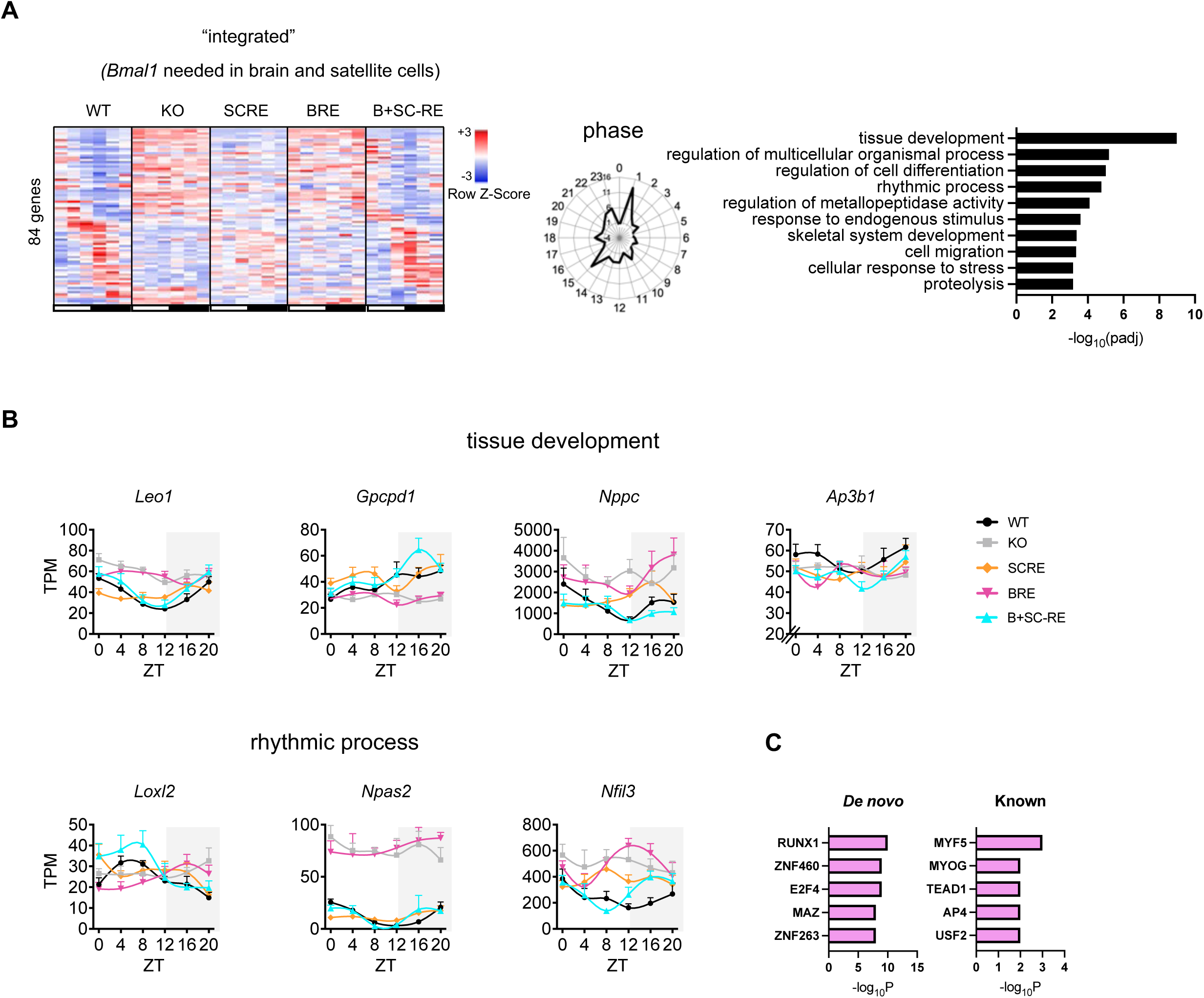
Integrated genes. **A**. Heatmap and phase-plot from dryR representing common oscillating genes in WT and B+SC-RE defined as integrated, and pathway enrichment from MSigDB (BICW>0.4, Amp>0.25,Cooks<1). Numbers on the perimeter of phase plots refer to ZT, numbers on internal grid to the number of oscillatory transcripts. **B**. Examples of oscillating genes belonging to pathways found in the enrichment analysis (KEA upregulated). TPM-Transcripts Per Million mapped reads, ZT-Zeitgeber Time. Wild-type (WT), Bmal1 total knock-out (KO), Bmal1 reconstitution in satellite cell only (SCRE), Bmal1 reconstitution in brain only (BRE), Bmal1 reconstitution in brain and satellite cells (B+SC-RE)**. C**. Transcription factors predicted by HOMER from the integrated.

**Figure S7.**
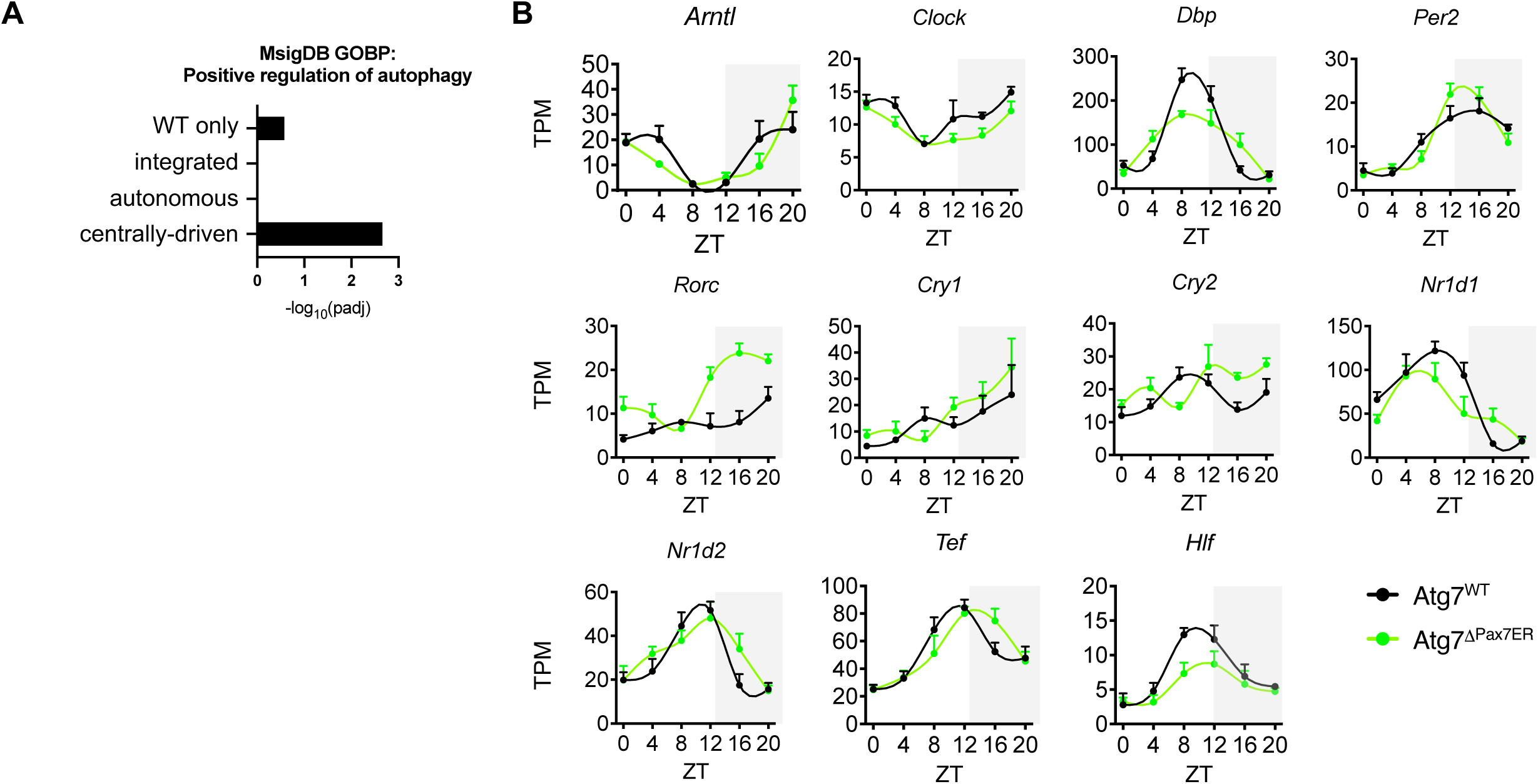
Autophagy and circadian rhythms. **A**. Enrichment from MSigDB for positive regulation of autophagy in selected categories. **B**. Core clock genes oscillations in Atg7^WT^ and Atg7^ΔPax7CreER^ satellite cells (BICW>0.6, Amp>0.25,Cooks<1). TPM-Transcripts Per Million mapped reads, ZT-Zeitgeber Time.

**Figure S8.**
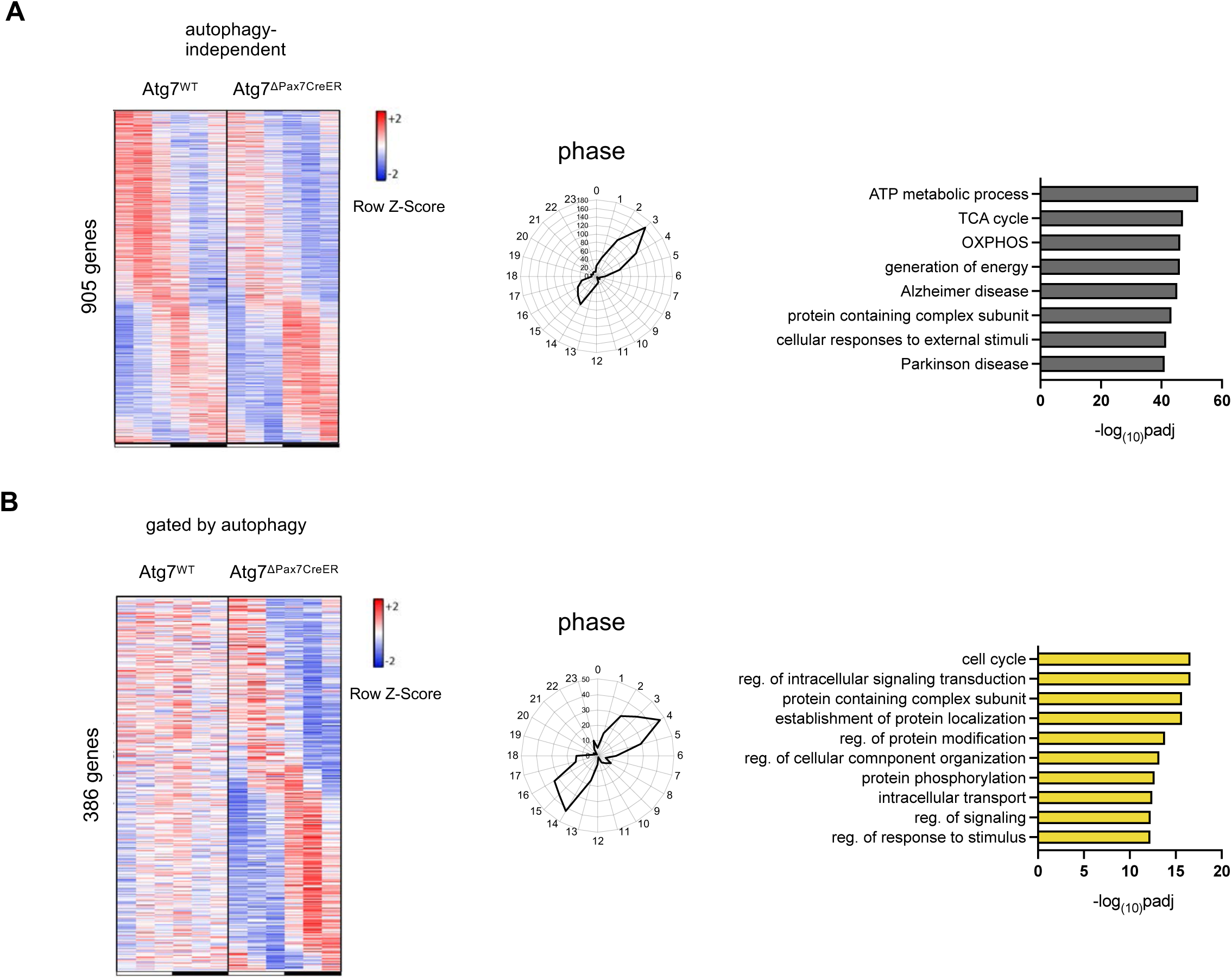
Autophagy gates oscillations of subsets of genes. **A**. Heatmap, phase plot and MSigDB pathway enrichment from a dryR model showing 905 genes whose oscillations do not depend on autophagy (BICW>0.6, Amp>0.25, Cooks <1). Numbers on the perimeter of phase plots refer to ZT, numbers on internal grid to the number of oscillatory transcripts (also applies to phase plot in B). **B**. Heatmap, phaseplot and pathway enrichment from a dryR model showing genes that gain oscillations in absence of autophagy.

